# Using simulation to aid trial design: ring-vaccination trials

**DOI:** 10.1101/071498

**Authors:** Matt D.T. Hitchings, Rebecca F. Grais, Marc Lipsitch

## Abstract

**Background:** The 2014-5 West African Ebola epidemic highlights the need for rigorous, rapid clinical trials under difficult circumstances. Challenges include temporally and spatially patchy transmission, and the responsibility to deliver public health interventions during a randomized trial. An innovative design such as ring vaccination with an immediate arm and a delayed arm can address these issues, but complex trials raise complex analysis issues.

**Methods and Findings:** We present a stochastic, compartmental model for a ring vaccination trial of a vaccine for an Ebola-like disease. After identification of an index case, a ring of primary contacts is recruited and either vaccinated immediately or after a delay of 21 days. The primary outcome of the trial is effectiveness calculated from cumulative incidence in the two arms, counting cases only from a pre-specified window in which the immediate arm is assumed to be fully protected and the delayed arm is not protected. The results of simulating the trial are used to calculate the sample size necessary for 80% power and the estimates of effectiveness are reported under a variety of assumptions regarding the trial design and implementation.

The three key components of sample size calculations – attack rate in controls, estimate of incidence difference between the arms, and intracluster correlation coefficient – are dependent on trial design and implementation in a way that can be quantitatively predicted by the model. Under baseline parameter assumptions, we found that a total of 8,900 study participants were needed to achieve 80% power to detect a difference in attack rate between the two arms, whereas a standard approach with the same parameters returns a necessary sample size of 7,100 individuals. Such a study would on average return a vaccine effectiveness estimate of 69.81%, with average 95% confidence interval (41.2%, 84.2%).

We found that for this design the necessary sample size and estimated effectiveness are sensitive to properties of the vaccine – in particular, pre-exposure and post-exposure efficacy; to two setting-specific parameters over which investigators have little control – rate of infections from outside the ring and overall attack rate in the controls; and to three parameters that are determined by the study design – the time window in which cases are counted, intensity of case-detection and administrative delay in vaccinating individuals.

This approach replaces assumptions about parameters in the trial with assumptions about disease dynamics and vaccine characteristics at the individual level.

**Conclusions:** Incorporating simulation into the trial design process can improve robustness of sample size calculations. Simulation can identify optimal values for study design parameters that can be controlled. For this specific trial design, vaccine effectiveness depends on properties of the ring vaccination design and on the measurement window, as well as the epidemiologic setting. Rejecting the null likely indicates one or more types of vaccine efficacy at the individual level, but the magnitude of the effect will vary across settings.

## 1. Introduction

The West African Ebola epidemic highlighted the need to identify a range of trial designs to evaluate vaccine effects rapidly, efficiently and rigorously during emerging disease outbreaks. The ring-vaccination trial approach employed in the *Ebola ça suffit* trial in Guinea is one innovative approach (1), which produced valuable evidence that the vaccine could prevent Ebola infection (2) and may become a paradigm for evaluation of future vaccines against emerging diseases. Other approaches considered include individual randomization and a stepped-wedge design.(3,4) Therefore in considering such a design in the future,it will be important to define the circumstances under which it is likely to be successful, so that these different study designs may be compared for their prospects of obtaining valuable data. Two key determinants of the value of the data produced by a particular design are (i) the sample size required to obtain power to test for vaccine effectiveness, and (ii) the likely point estimate of effectiveness obtained by the trial design, for a given set of conditions. We use a simulation model of a ring-vaccination trial to evaluate these two aspects under various assumptions about the properties of the vaccine, the trial, and the population.

Although the only implementation of the ring trial design has been in Guinea during the Ebola epidemic, lessons can be learned and extended to other diseases and contexts. Here, we examine the tail end of an epidemic of a disease with a latent and asymptomatic phase with effective contact tracing to illustrate a more widely-applicable set of findings. In particular, we use baseline parameters values consistent with Ebola in West Africa in 2014-5, but we vary several assumptions over broader ranges than those occurring in the *Ebola ça suffit* trial, with the aim of being relevant to a range of potential future situations.

## 2. Methods

### 2.1. Ring vaccination trial

The simulation is based on a stochastic, susceptible-exposed-infectious-detected-removed-vaccinated (SEIDRV) model for individual disease events, and it represents progression of the disease within the context of a ring vaccination trial, wherein a ring is randomized either to immediate vaccination (on day 1) or delayed vaccination (on day 22), and all individuals in the ring receive vaccination on that day.(2) Thus in the baseline scenario we assumed that no individuals are ineligible or refuse the vaccine, so that all susceptible or exposed individuals in the ring are vaccinated, and that there is no heterogeneity or administrative delay affecting the day of vaccination. When we vary the administrative delay, we assume that the delay applies to both arms, so that the delayed arm is still vaccinated 21 days after the immediate arm.

Once a ring in the trial is initiated by first identifying an index case, there are 50 contacts who make up that index case’s ring. There is no spillover between rings, and the index case is not included in the estimation of vaccine efficacy. A ring is counted whether or not it contains any cases infected by the index case or by external infection; some rings will have secondary attack rates of 0 because the index was detected and isolated before he infected anyone else, and no external infection occurred. During the follow-up of a ring, each case is detected at with daily probability pH.

Because both arms receive the vaccine, cases are only counted during a window in which the immediate arm is presumed to be protected by the vaccine, and the delayed arm is not protected. The window length is set to 21 days, equal to the vaccination delay between the arms, and the start of the window is set at baseline to be 16 days from index case identification, which is a natural choice as the sum of the vaccine ramp-up period and the mean incubation period.(1) Once detected, secondary cases are counted towards the cumulative incidence in an arm if they developed symptoms during the time window (though they may be detected after that window ends).

For additional details on the disease transmission model, ring initiation, and analysis of the trial see the supplementary appendix.

### 2.2. Choice of parameters

Table 1 shows the parameters used in the model, their meanings, values under baseline assumptions, and references or justifications.

**Table 1:**
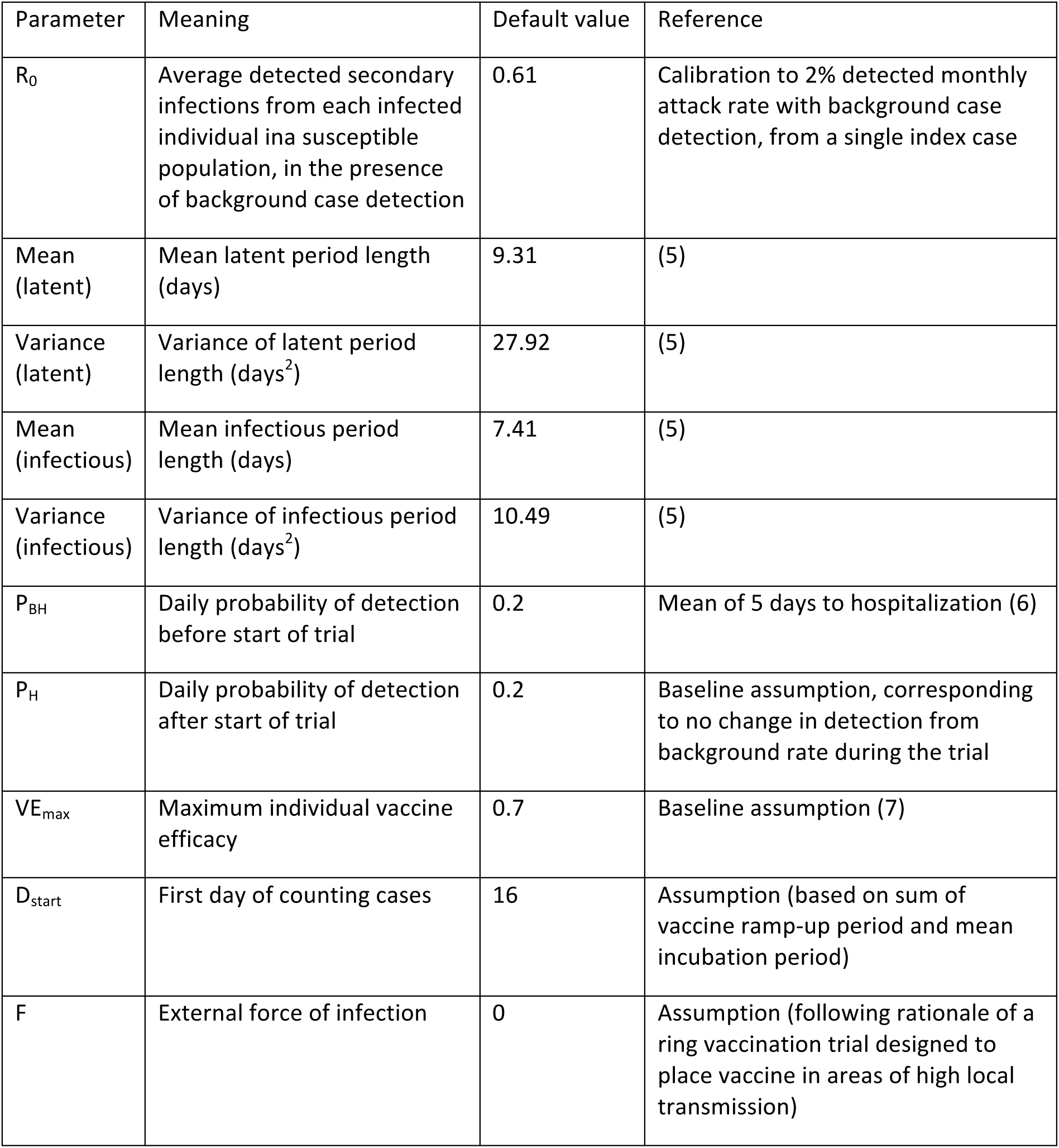
Table of parameter values and meanings, and references for those parameters which were chosen using the literature

| Parameter | Meaning | Default value | Reference |
| --- | --- | --- | --- |
| $R_0$ | Average detected secondary infections from each infected individual in a susceptible population, in the presence of background case detection | 0.61 | Calibration to 2% detected monthly attack rate with background case detection, from a single index case |
| Mean (latent) | Mean latent period length (days) | 9.31 | (5) |
| Variance (latent) | Variance of latent period length (days<sup>2</sup>) | 27.92 | (5) |
| Mean (infectious) | Mean infectious period length (days) | 7.41 | (5) |
| Variance (infectious) | Variance of infectious period length (days<sup>2</sup>) | 10.49 | (5) |
| $P_{BH}$ | Daily probability of detection before start of trial | 0.2 | Mean of 5 days to hospitalization (6) |
| $P_H$ | Daily probability of detection after start of trial | 0.2 | Baseline assumption, corresponding to no change in detection from background rate during the trial |
| $VE_{max}$ | Maximum individual vaccine efficacy | 0.7 | Baseline assumption (7) |
| $D_{start}$ | First day of counting cases | 16 | Assumption (based on sum of vaccine ramp-up period and mean incubation period) |
| $F$ | External force of infection | 0 | Assumption (following rationale of a ring vaccination trial designed to place vaccine in areas of high local transmission) |

The context we are modeling is the end of an epidemic, so that R_0_ is less than one. In order to calibrate the model, we set R_0_ to reproduce a monthly detected attack rate of 2% when starting from one infected individual in a ring of 50 susceptible individuals, in the presence of background case detection and the absence of vaccination.

## 3. Results

Under the baseline parameter assumptions listed above, the sample size necessary in each arm to achieve 80% power to detect a difference in cumulative incidence between the two arms is 89 rings, each containing 50 individuals, making a total of 8,900 study participants. This trial would on average return a vaccine efficacy estimate of 69.81%, with average 95% CI (41.2, 84.2).

### 3.1. Determinants of vaccine effectiveness estimate

Under baseline parameters in this model, the median vaccine effectiveness calculated from performing 100 trials with 89 rings in each arm was 70%. This value should include direct and indirect effects, so we would expect it to exceed the direct effect of 70%. However, while direct effects begin immediately, indirect effects are only important in the second generation of preventable cases onwards. The time window was chosen to capture a period of time in which the immediate arm receives full protection of the vaccine and the delayed arm receives none. The length of the window (21 days) is slightly larger than the average generation interval (17 days) so on average it will not capture many members of this second generation in the immediate arm. If we were to extend the time window to capture indirect effects, we would include time in which the delayed arm is receiving protection from the vaccine, leading to a downward bias in the vaccine effectiveness estimate.

Figure 1 shows the effect of six variables on the point estimate of vaccine effectiveness: daily probability of detection, true individual vaccine efficacy, force of external infection, baseline attack rate in the unvaccinated population, administrative delay in vaccination, and start day of case-counting window.

**Figure 1:**
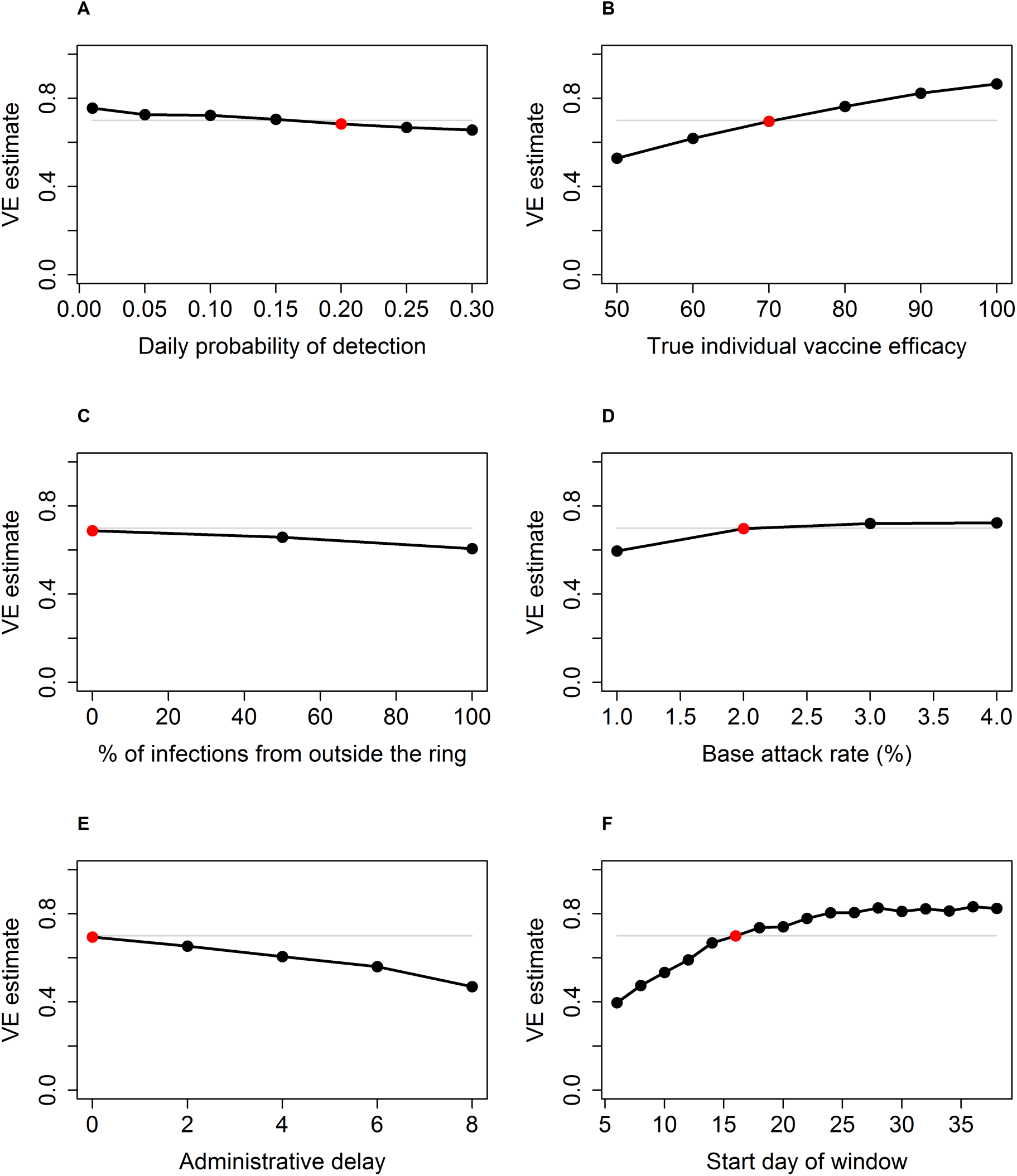
Point estimate of vaccine effectiveness derived from a trial with 80% power to detect vaccine effect shown against: (left to right, top to bottom) A: daily probability of detection, B: true individual vaccine efficacy, C: force of external infection, D: baseline attack rate in the unvaccinated population, E: administrative delay in vaccination, and F: start day of case-counting window. In each panel, the VE estimate corresponding to the baseline parameter set is highlighted in red, and the gray line represents the individual vaccine efficacy of 70%. All other parameters are set at the baseline values.

Firstly, if there is enhanced surveillance in both arms of the trial leading to more rapid isolation of infectious cases (p_H_>p_BH_), this will modestly reduce effectiveness estimates (Figure 1A). Secondly, the properties of the vaccine and the immune response in the individual (encapsulated in the model by maximum efficacy, ramp-up time, and post-exposure efficacy) work together to determine the effectiveness in a population in a way that we cannot express in a closed form. Qualitatively, as individual efficacy increases the effectiveness will increase (Figure 1B). Vaccine efficacy estimate also increases with increasing post-exposure efficacy (Figure S1). Thirdly, the percentage of infections from within the ring shows a weak negative association with the estimate of vaccine effectiveness (Figure 1C). While the magnitude of indirect effects is modest as discussed above, they are almost negligible when most infections are from outside the ring, because preventing infections within the ring does not confer as much protection to susceptible individuals. Delay between ring formation and vaccination means that by the beginning of the time window the vaccine has had less time to prevent cases in the immediate arm. Thus the reduction in incidence in the immediate arm does not reflect the true effect of the vaccine and the vaccine effectiveness estimate is reduced (Figure 1D). A higher attack rate leads to more cases in both arms, so in the immediate arm there will be a modest increase in tertiary cases (i.e. those that could be prevented by the indirect effect of the vaccine) and thus a small increase in indirect effects (Figure 1E).

A major determinant of the effectiveness estimate is the choice of time window in which to count cases, as seen in Figure 1F. Not surprisingly, starting the window too early reduces the estimated effectiveness because it includes a period of time during which the vaccine cannot affect the incidence of cases becoming symptomatic – many cases becoming symptomatic on day 8, for example, will have been infected by the index case prior to isolation, or will have been infected by a contact on (say) day 3, before the vaccine had time to induce protection.

Starting the window later than the baseline of 16 days allows the trial to capture later generations in the chain of transmission, from a vaccinated person to another vaccinated person. This increases the vaccine effectiveness estimate as it includes indirect effects. One might expect to see that starting the window too late would reduce effectiveness estimates because it would include a period when the delayed group was also protected by the vaccine. This does not appear to be the case, at least up to a start time of 35 days (Figure 1F). Before vaccination, incidence in both arms is decreasing at the same exponential rate, and thus in proportion to each other. The effect of vaccination is to increase the rate of decline in the immediate arm by interrupting potential transmission chains. The difference between the two arms increases as indirect effects come into play, until the delayed arm receives vaccination. The effect of vaccination in the delayed arm is to increase the rate of decline so that it is equal to the rate in the immediate arm. This explains why the VE estimate doesn’t decrease for later time windows; the incidence in the delayed arm doesn’t ‘catch up’ with that in the immediate arm, it merely ‘keeps pace’ when the vaccine begins to have an effect. Figure 2 shows, on the log scale, the change in incidence rate decline in the delayed arm that happens around day 30-35, or 9-14 days after vaccination. After that, the two lines are parallel on the log scale, meaning that they are declining in proportion and so the VE estimate, which is based on the incidence ratio, doesn’t change.

**Figure 2:**
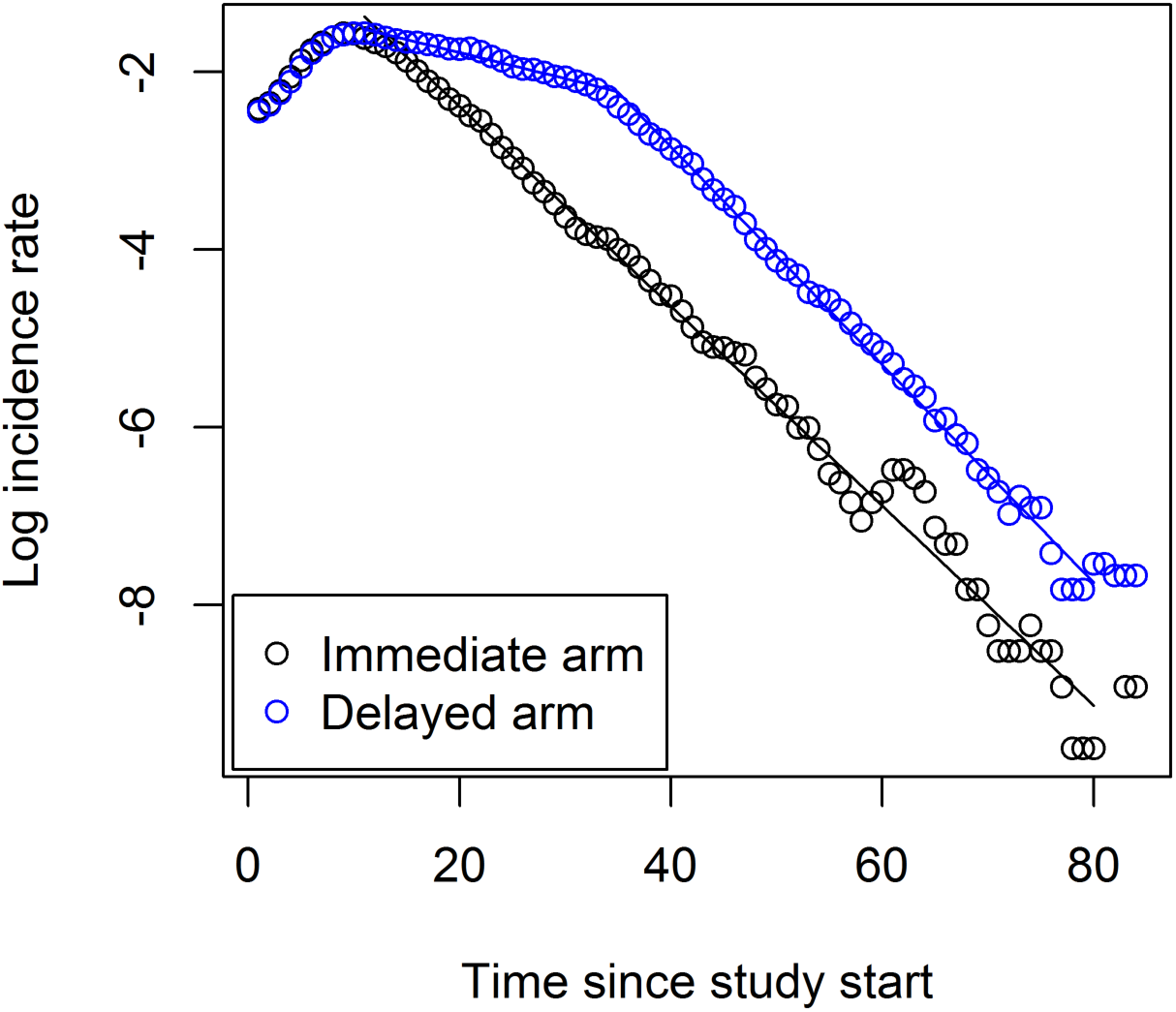
Simulated log incidence rate of detected disease in the trial, in the immediate arm (black circles) and delayed arm (blue circles), with linear fit in the immediate arm (black line) and piecewise linear fit in the delayed arm (blue line). The change in rate in the delayed arm corresponds to the direct effect of the vaccine. Circles represent means over 15,000 simulations.

### 3.2. Determinants of sample size

Figure 3 shows the effect of six variables on the required sample size: baseline attack rate in unvaccinated population, start day of case-counting window, daily probability of detection, true individual vaccine efficacy, administrative delay in vaccination, and force of external infection.

**Figure 3:**
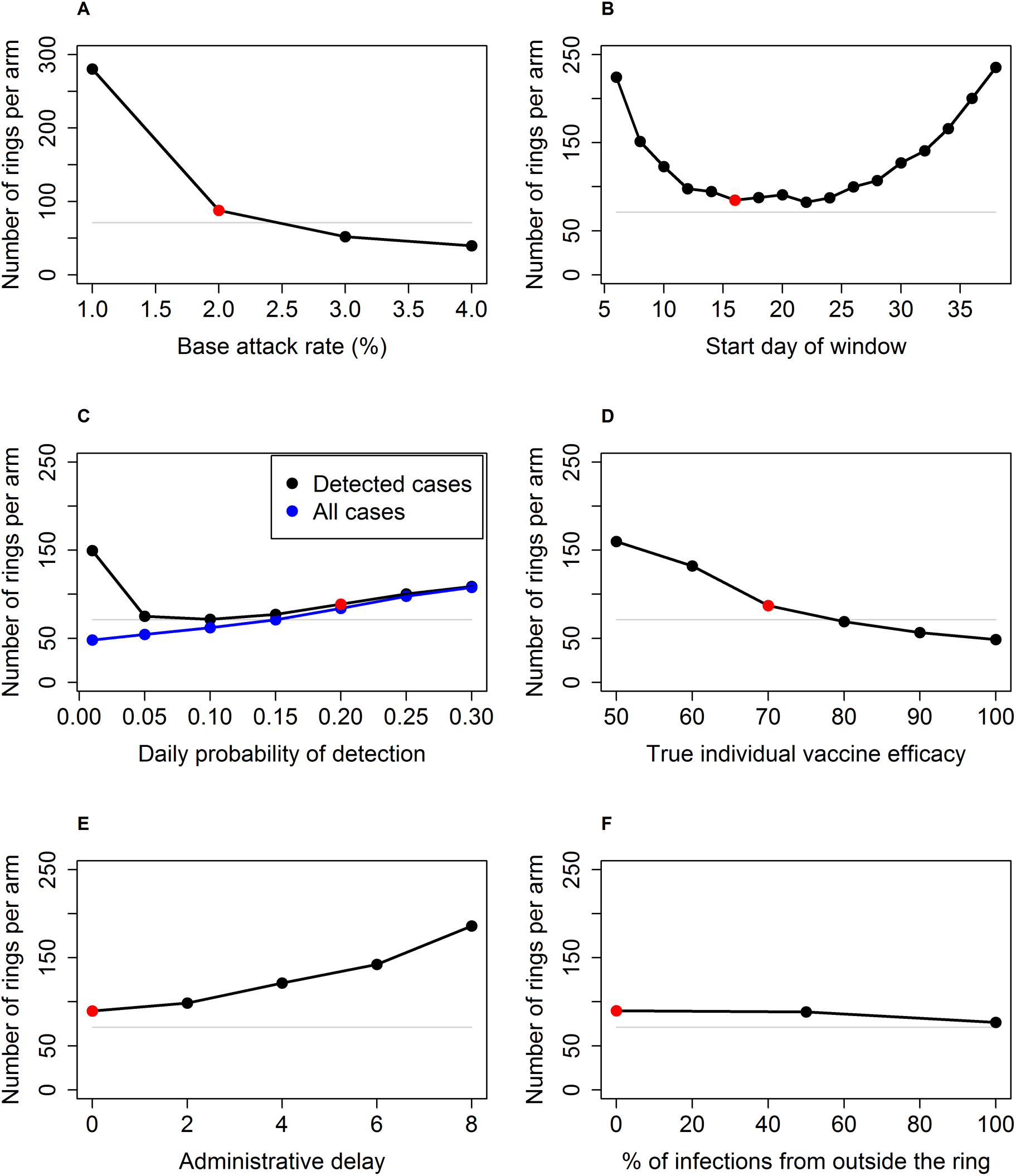
Number of rings per arm required to achieve 80% power to detect a difference in cumulative incidence between the two arms against six key parameters: (left to right, top to bottom) A: baseline attack rate in unvaccinated population, B: start day of case-counting window, C: daily probability of detection, D: true individual vaccine efficacy, E: administrative delay in vaccination, and F: force of external infection. In Figure 3C, sample sizes are shown for VE estimates based on only detected cases (black) and on all cases (blue). In each panel, the sample size estimate corresponding to the baseline parameter set is highlighted in red, and the sample size derived from a standard approach is shown by the gray line. All other parameters are set at the default values.

The effect of each parameter on the sample size can be understood through its effect on one or more of the three factors that determine the power of this trial: the number of events, how they are distributed between the two arms, and the level of clustering. Respectively these factors are represented by the attack rate in the controls, the cumulative incidence difference between the arms, and the intracluster correlation coefficient (ICC).(8)

Variables that decrease the incidence rate in the controls and cases will decrease the power because for the same sample size the trial will observe fewer events. The baseline detected attack rate among unvaccinated individuals is a simple example of such a parameter (Figure 3A). Two other parameters act on the overall incidence in the trial. Firstly, making the start of the case-counting window later decreases incidence in both arms because with R_0_<1 the incidence is on average declining (Figure 2), so across all rings in the trial the number of cases decreases over the follow-up period (Figure 3B). Secondly, the case detection decreases detected incidence rate at both extremes (Figure 3C). When case detection is high, transmission chains are interrupted by case isolation and the true incidence decreases. When case detection is low, many cases die or recover before they can be detected and consequently the detected incidence decreases.

Variables that make the two arms of the trial appear more different will increase the power of the trial since the ability to differentiate between them is increased, and Figure 1 identifies such variables. Vaccine characteristics, in particular vaccine efficacy (Figure 3D), are simple examples of such a parameter, since the immediate arm receives greater protection against disease compared to the delayed arm. Changes to two other parameters increase the incidence difference in this way, as explained above: reducing the delay between ring formation and vaccination (Figure 3E) and starting the case-counting window earlier (Figure 3B).

The effect of the timing of starting to count cases thus reflects two opposing forces on the sample size: it decreases sample size by increasing the incidence difference, and it increases sample size by decreasing the overall incidence. When the window is early, the former of these effects dominates as seen by the increase in sample size for early time windows in Figure 3B. When the window is late, the latter effect dominates, as seen by the increase in sample size for late time windows in the same figure.

Finally, the level of clustering in the population inflates the sample size. It is often not intuitive to predict the direction in which a parameter will cause the ICC to change, and in many cases the ICC is not sensitive to the parameter. One exception is the infection from outside the ring (Figure 3F). The most significant effect of introducing external infection and reducing within-ring transmission is to decouple infection probability for one individual within a ring from that of others within the same ring. This reduces clustering in incidence (making it more Poisson-like), thus reducing the ICC and the necessary sample size.

## 4. Discussion

The ring-vaccination, cluster-randomized design has two key strengths that make it a good candidate when disease transmission exhibits spatiotemporal variation. Firstly, by including members of the study population who are contacts of cases, the trial preferentially selects those at higher risk of disease acquisition, leading to an increase in efficiency while preserving Type I error through randomization. Secondly, even those study subjects who are randomized to delayed vaccination are theoretically in close contact with the study team meaning that the population who are at the highest risk are followed closely.(9)

In addition, vaccination of clusters when they arise allows for roll-out of the vaccine, meaning that this design is appropriate when logistical constraints make immediate mass vaccination impossible or inappropriate. In this respect it is similar to a stepped-wedge cluster trial, in which prespecified clusters within the study population are vaccinated in a random order. Although we have not made a direct comparison in this study, Bellan et al (10) showed that the stepped-wedge design is underpowered when the incidence declining because it cannot prioritize the vaccine for those at highest risk. The ring vaccination design, on the other hand, is inherently risk-prioritized because all study participants should be at higher risk than the general population.

All trials should be correctly powered in order to avoid erroneous rejection of an efficacious vaccine, leading to a waste of valuable resources. For a trial design with several complexities such as the one presented here, a sophisticated approach to sample size calculation is merited. A standard approach to sample size calculation for this trial would involve specifying the attack rate among the controls, the desired effect of the vaccine on the population level, and the ICC. In the context of a serious epidemic, these parameters are unlikely to be estimated with certainty; for example, the ICC requires cluster-level data to be estimated accurately. Therefore, the modeling approach replaces assumptions about these cluster-level quantities with assumptions about population-level parameters and disease characteristics, which are more likely to be available through analysis of data from the outbreak.

A second advantage of the modeling approach is that, based as it is on a simulating the transmission of disease within a trial, it is possible to explore the impact of parameters describing the design of the trial and the properties of the disease. The added detail gained from specifying the disease model allowed us in this study to identify some key issues with the design that are worth considering.

Firstly, as seen in Figure 3C, increasing case-finding efficiency above background rate has a negative impact on power, as fast isolation of cases in both arms leads to an overall decrease in cases observed by the trial. In future trials it is worth considering if there are alternative endpoints that can be used that could allow for less intensive follow-up or post-trial detection of cases.

Secondly, a key design consideration in the delayed-arm ring-vaccination trial is when to count cases. An intuitively appealing approach is to place the window so that the immediate arm is receiving full protection and the delayed arm none. This should in theory minimize bias caused by misclassification of unvaccinated individuals as vaccinated and vice-versa. While this placement achieves nearly maximal power, it does not maximize the VE estimate. Indirect effects that are important later in time increase the VE estimate for later time windows, while at the same time declining incidence within each ring decreases power for later time windows.

Finally, the above point draws attention to the fact that caution is required when interpreting the VE estimate produced by the trial. As seen in Figure 1, many parameters that are not characteristics of the vaccine can influence the estimated effectiveness. Whether this is due to misclassification (for example, when the time window is too early) or due to indirect effects (for example, when the attack rate is high enough to measure significant protection of tertiary cases), the context of the trial should be taken into account when considering the VE estimate.

The focus of this model was to explore parameters within each ring and understand how they affect the quality of data coming from the trial. As a result, we did not consider the wider context of the population disease dynamics, and in particular how and when the rings arise. For example, we calibrated R_0_ to a secondary attack rate in a cluster was 2%, which is not necessarily comparable to the monthly cumulative incidence in the population. If transmission takes place mainly in clusters then population cumulative incidence could be somewhat lower than cluster secondary attack rate, increasing the efficiency of a ring-vaccination trial relative to a stepped-wedge cluster trial or individual RCT. Linking this model to a model of disease within the general population would allow us to make direct comparisons to other trial designs such as the stepped-wedge cluster trial and the individually-randomized trial investigated elsewhere,(10, 11) but it would require detailed 13 information about the nature of clustering of the disease in this context, and for simplicity we focused on the within-ring dynamics only.

As with every model, there are limitations to these simulation results. The strength of the modeling approach compared with a standard approach is that it better estimates the parameters on which the sample size depends. However, some of the model parameters might still be uncertain in a situation in which such a model might be useful. For example, we are likely to have little information about the characteristics of a disease, in particular its latent and incubation period, and its R_0_. The simulation results are heavily dependent on these assumptions, and so they cannot be used at the very outset of epidemic, or else they risk being highly inaccurate. Even at the end of the West African Ebola epidemic, there were no more than four or five reliable calculations of the latent and infectious periods of EVD, and indeed there is perhaps evidence that the latent and incubation periods do not precisely overlap.(12) In addition, we have considered only the simplest method of analysis for the trial – a comparison of attack rates between the two arms after correction for clustering of cases within rings. More sophisticated methods, including time-to-event analyses incorporating ring-level random effects, as performed in the *Ebola ça suffit* trial, would have somewhat different sample size requirements.

In an epidemic situation, when the R_0_ has changed over time and with a previously untested vaccine, power calculations can be very sensitive to parameters about which very little is known. Simulations such as these can be important aids in understanding a range of values for these parameters before a trial is carried out, and thus ensuring that the trial has sufficient power to detect an efficacious vaccine. In this trial, a finding significantly different from the null likely indicates one or more types of vaccine efficacy at the individual level, but the magnitude of the effect and the power to detect the effect will vary across settings.

## Supplementary Information

S1 Appendix. Additional detail on methods, including disease transmission model and simulation and analysis of trial.

## S1. Appendix

### S1.1. Disease transmission model

To simulate the spread of a disease within a small (m=50) community of individuals who have close contact with each other (henceforth a “ring”), we used a stochastic, compartmental model with five compartments: susceptible, exposed, infectious, isolated, and removed (either recovered or dead). The time step was one day, and all processes such as infection, disease progression, etc. are discretized to occur at the end of a particular day. We assumed that individuals become infectious when symptoms appear, meaning that the latent period and incubation period are concurrent (WHO. http://www.who.int/mediacentre/factsheets/fs103/en/). Each individual in the susceptible compartment has a daily force of infection from two sources: externally from individuals not contained within the ring, and internally from individuals within the ring. The former is denoted by a fixed hazard F/day, and the latter has hazard equal to βI/day, where β is the transmission rate constant and I is the number of infectious individuals in the ring. An individual who becomes infected is placed in the exposed compartment, where they spend a number of days determined by a gamma distribution, with mean 9.31 days and variance 27.92 (days^2^).[1] At the end of the latent period, the individual is moved into the infectious compartment, where they spend a number of days determined by an independent gamma distribution with mean 7.41 days and variance 10.49 (days^2^).[1] While an individual is in the infectious compartment, they have a per-day probability of being detected and isolated, p_H_. The act of isolation immediately ends their infectiousness, meaning that case detection stops transmission. This phenomenon is a departure from how the *Ebola ça suffit* trial was run, but we model it here because it introduces interesting design issues. If they reach the end of the infectious period without being detected, they are placed into the removed category, at which point they are no longer infectious. In this model we have not allowed for a *post mortem* period of infectiousness, nor the possibility of sexual transmission among those recovered, nor of asymptomatic infections. Finally, we assume that the epidemic is ending due to behavior change and other factors, rather than depletion of susceptibles. Therefore there are no recovered individuals in the initial population.

### S1.2. Ring vaccination trial details

Initially, we considered a vaccine whose only effect was pre-exposure prophylaxis; in initial runs we assumed the vaccine had no effect if given to a person who was exposed but not yet infectious, an assumption we later relaxed. After administration of the vaccine, a ramp-up period (Dramp, set in the baseline runs to 7 days) occurs during which vaccine efficacy (VE) rises linearly from 0% to the maximum individual efficacy (VEmax, set at baseline to 70%), after which there is no change in the individual efficacy over the study period.[2] The vaccine is leaky,[3] providing partial protection to all individuals, so that the effect of the vaccine is to reduce the force of infection on a vaccinated individual by a factor of (1–VE). When included, post-exposure vaccine effects were modeled as 15 follows: when a vaccinated, latently infected subject leaves the exposed class, he will move straight to the removed class with probability p_PEP_.

To initiate rings, we simulated the following steps: One infected individual (not counted in the m=50) is infected, and the length of his infectious period is drawn from the gamma distribution. As he progresses through his infectious period, susceptible members of the ring can be infected with daily probability 1 – e^−βI+F^. In addition, the index case can be detected and isolated with daily probability p_BH_. For baseline simulations we set p_BH_=0.2, meaning that it takes on average 5 days to detect and isolate an infected individual.[4] If he is detected before his infectious period ends, he is rendered non-infectious by isolation, and he becomes the index case for a ring, which proceeds as described below. If he is not detected before recovering or dying, he is effectively invisible to trial investigators, so he is not counted as an index case, and the simulation is terminated and repeated again, without counting the “invisible” case as part of the study sample.

### S1.3. Trial simulation and analysis

To calculate the sample size required for a desired power, a cluster-level analysis is performed, using a t-test to compare cumulative incidence [5] in the two arms from a simulated trial with k=15,000 rings. This sample size calculation includes an inflation factor (1+ρ*(m-1)), whereρ is the intracluster correlation coefficient (ICC), calculated as per Shoukri,[6] adjusting for the covariate of trial arm. The cumulative incidence of detected EVD cases is recorded in each arm of the trial, and the vaccine effectiveness is estimated as VE_est_=(1 – CI_imm_/CI_del_)*100, where CI_imm_ is the cumulative incidence in the immediate arm and CI_del_ is the cumulative incidence in the delayed arm. Since we expect the event to be rare, this calculation of vaccine effectiveness will be approximately equal to the measure using the hazard ratio.[3] As we are assuming no vaccine ineligibility or refusal, this quantity estimates the combination of the direct and indirect effect of the vaccine.[7] In order to output the likely estimate of vaccine effectiveness derived from this trial, we perform the trial 100 times at the required sample size calculated above, and we report the median vaccine effectiveness estimate from these 100 trials. All simulation was performed using R. [8]

---

**S1 Figure.**
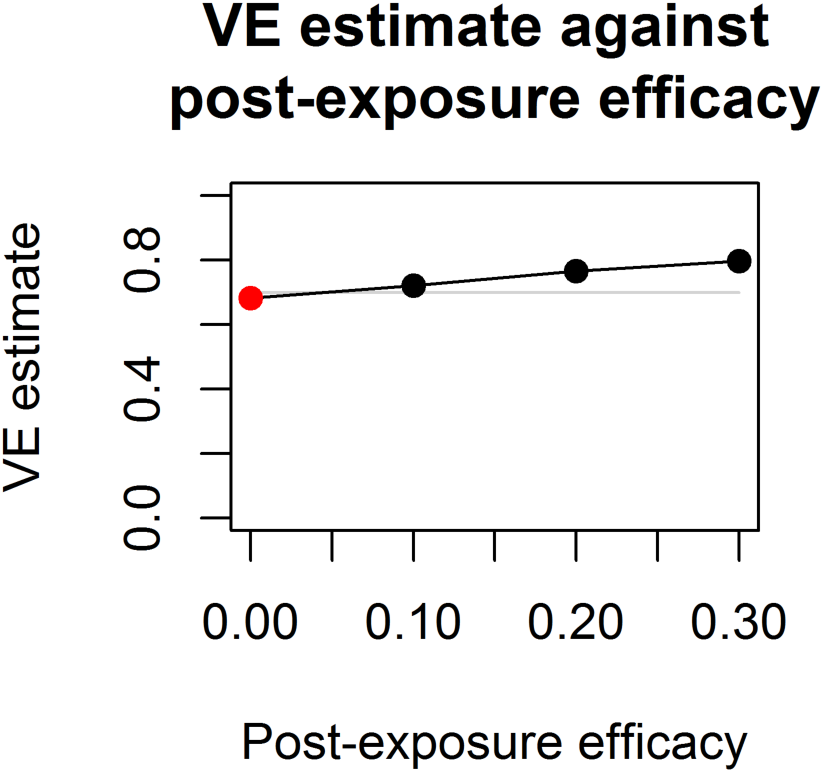
Point estimate of vaccine effectiveness derived from a trial with 80% power to detect vaccine effect shown against post-exposure vaccine efficacy.

## S1. R code used in this study

~~~
#Updated July 28 2016 by Matt Hitchings
#By Marc Lipsitch, Harvard T.H. Chan School of Public Health
# Ring vaccine simulation comparing two ring delays
# In this update I have rewritten the main loop (i.e. the simulation of the trial) as a function, to
# make it more flexible. This way I can, in a separate script, run the trial once to determine
# the required sample size, then run it again multiple times to get the vE CI.
# Results should not be shown publicly without written permission of Marc Lipsitch
mlipsitc@hsph.harvard.edu
simulatetrial<-function(pdhosp, VEmax, startcount,nsim,R0,extfoi_numerator, admin_delay=0, VEtake=6, PEPVE=0, Vuptake=1, window_length=21) {
 #Defaults for inputs: pdhosp=0.16, VEmax=0.7, startcount=18, R0=0.61, extfoi_numerator=0, VEtake=6, PEPVE=0

 # Changeable parameters
 pdhosppre<-0.2
 startcount<-startcount+admin_delay
 endcount<-startcount+window_length

 #FIXED SIMULATION PARAMETERS
 m<-50 #ring size
 endfu<-84 #end of followup 12 weeks
 tvacs =admin_delay+c(1,1+window_length) #day vaccine introduced

 #natural history
 maxnE<-21 #maximum days incubation/latent
 maxnI<-14 #maximum days infectious
 meannI<-7.41 #mean days infectious, used to calculate beta
 nSV<-endfu # days post vaccine to track
 people’s susceptibility
 extfoi<-extfoi_numerator/100/endfu #foi per capita from outside the ring - default to 1%
over the full course of observation (so divide by endfu)
 beta<-R0/m/2.94# daily infectiousness of an I
 nSV<-endfu # days post vaccine to track people’s susceptibility

 # Parameters for gamma distribution of incubation and infectious periods
 inc_shape<-3.04
 inc_rate<-0.33
 inf_shape<-5.29
 inf_rate<-0.71
 # Make vectors of conditional probability for whether individuals go to
 # next stage of infection
 # I’ve specified the maximum length of each period, so the last entry of the
 # vector is 1
 inc_probs<-c(rep(0,maxnE-1),1)
 inf_probs<-c(rep(0,maxnI-1),1)
 for (index in 1:(maxnE-1)) {
  inc_probs[index]<-(pgamma(index,inc_shape,inc_rate) - pgamma(index-
1,inc_shape,inc_rate))/(1-pgamma(index-1,inc_shape,inc_rate))
 }
 for (index in 1:(maxnI-1)) {
 inf_probs[index]<-(pgamma(index,inf_shape,inf_rate) - pgamma(index-
1,inf_shape,inf_rate))/(1-pgamma(index-1,inf_shape,inf_rate))
 }

 # Vaccine efficacy
 #starts protecting on day VEstart, ramps up over VEtake days to VEmax

 VEstart<-1
 VES<-c(rep(0,VEstart),(VEmax/VEtake)*(1:VEtake), rep(VEmax,endfu-VEstart-VEtake)) #Vaccine
efficacy vs infection each day after vaccination
 VEI<-0 #reduced infectiousness if any of vaccinated person who becomes infected (before or
after vaccination)
 betav<-beta*(1-VEI)
 SAR<-vector()
 trueSAR<-vector()
 pi<-vector()

 library(Hmisc)

 # A function that takes a vector and outputs the indices of the non-zero
 # elements, with duplicates for elements that are greater than 1
 vectortoindices<-function(vec) {
  indices<-which(vec>0)
  output<-vector()

  for (index in indices) {
   for (count in 1:vec[index]) {
     output<-c(output,index)
    }
  }
  output
 }

 # A function to do multiple outcomes from a function
 list <- structure(NA,class="result")
 "[<-.result" <- function(x,...,value) {
  args <- as.list(match.call())
  args <- args[-c(1:2,length(args))]
  length(value) <- length(args)
  for(i in seq(along=args)) {
    a <- args[[i]]
    if(!missing(a)) eval.parent(substitute(a <- v,list(a=a,v=value[[i]])))
  }
  x
 }

 # CORE FUNCTIONS
 #A time-advancing function to slide people along the conveyor,
 # possibly switching from end of state OLD to beg of state NEW
 tplus<-function(old,new){
  c(tail(old,1),head(new,-1))
}

 # A function to randomly pick, for each exposed and infected individual,
 # whether they remain in that class or move to the next class.
 random_periods<-function(old,probabilities){

  ones_switching<-rbinom(length(old),old,probabilities)
  ones_staying<-old-ones_switching
  num_switching<-sum(ones_switching)

  period_lengths<-vector()
  if (num_switching>0){
    period_lengths<-vectortoindices(ones_switching)
  }
  list(out1=num_switching,out2=ones_staying,out3=period_lengths)
 }

 # A function to take the vector of hospitalized individuals and update a data
 # frame with info on when they were hospitalized and when they became infectious
 # This will only work if subjects can only be hospitalized during their
 # infectious period, which is a reasonable assumption
 hosptodayinf<-function(daysinfhosp,tohosp,t){
   # Each entry of tohosp represents the number of individuals hospitalized
   # on that day, which is the day of their infectious period. For each entry
   # make a new vector that contains the day of hospitalization (measured in
   # time since trial start) and the day of symptom onset
   for (index in 1:length(tohosp)) {
     if (tohosp[index]>0) {
      for (subject in 1:tohosp[index]){
        # Day of hospitalization is t, day of symptom onset is t-index+1
        temp<-data.frame(dayinf=t-index+1,dayhosp=t)
        daysinfhosp<-rbind(daysinfhosp,temp)
       }
     } else {}
  }

  daysinfhosp}
 # # advance time and progress state, possibly incorporating PEPVE
 update<-function(state,pdhosp)
 {
   x<-state

 # Advance unvaccinated exposed and infected, taking into account random incubation/
 # infectious periods
 list[incu_numswitch,incu_stay,inculengths]<-random_periods(x$EU,inc_probs)
 newEU<-tplus(0,incu_stay)
 list[infu_numswitch,infu_stay,infulengths]<-random_periods(x$IU,inf_probs)
 newIU<-tplus(incu_numswitch,infu_stay)
 newRU<-x$RU+infu_numswitch

 # Advance vaccinated exposed and infected, taking into account random incubation/
 # infectious periods
 list[incv_numswitch,incv_stay,incvlengths]<-random_periods(x$EV,inc_probs)
 newEV<-tplus(0,incv_stay)
 newSV<-tplus(0,x$SV) # vaccinated susceptibles move en masse to being vaccinated for one more day
  list[infv_numswitch,infv_stay,infvlengths]<-random_periods(x$IV,inf_probs)
  IVPrev<-rbinom(1,incv_numswitch,PEPVE) #prevention by PEP of progression
  # from last day of EV
  newIV<-c(incv_numswitch-IVPrev,head(infv_stay,-1)) #else progress to IV[1]
  newRV<-x$RV+infv_numswitch+IVPrev
  x$EU<-newEU
  x$IU<-newIU
  x$RU<-newRU
  x$SV<-newSV
  x$EV<-newEV
  x$IV<-newIV
  x$RV<-newRV

 # If anyone switched from I to R or E to R, record the day. This is so we can censor
people for survival analysis
 newRs <- infu_numswitch + infv_numswitch + IVPrev
 onestep_Rdays <- rep(time,newRs)

 # This is to record the lengths of incubation and infectious periods, both
 # as a check and as a potentially interesting outcome
 inc_lengths<-c(inculengths,incvlengths)
 inf_lengths<-c(infulengths,infvlengths)

 # Vaccinate - this section will work only if vaccination is on a fixed day tvac for
everyone, and happens only once.
 if (time==tvac){
   # Include vaccine refusal/ineligibility
   vacS<-rbinom(1,x$SU,Vuptake)
   x$SV<-c(vacS,rep(0,endfu-1))
   x$SU<-x$SU-vacS
   vacE<-rbinom(maxnE,x$EU,Vuptake)
   x$EV<-vacE
   x$EU<-x$EU-vacE
   vacI<-rbinom(maxnI,x$IU,Vuptake)
   x$IV<-vacI
   x$IU<-x$IU-vacI
   x$RV<-rbinom(1,x$RU,Vuptake)
   x$RU<-x$RU-x$RV
  } else{}
 # Infect
 foi<-extfoi+beta*sum(x$IU)+betav*sum(x$IV)

 incU<-rbinom(1,x$SU,1-exp(-foi))
 incV<-rbinom(nSV,x$SV,(1-exp(-foi))*(1-VES))
 x$EU[1]<-incU
 x$EV[1]<-sum(incV)
 x$SU<-x$SU-incU
 x$SV<-x$SV-incV

 #hospitalize - assume this ends their infectivity
 tohosp<-rbinom(maxnI,x$IU,pdhosp)
 x$IU<-x$IU-tohosp

 tohospv<-rbinom(maxnI,x$IV,pdhosp)
 x$IV<-x$IV-tohospv
 x$H<-x$H+sum(tohosp)+sum(tohospv)

 # Get info about day infected and hospitalized for those hospitalized
 tohosptot<-tohosp+tohospv
 daysinfhosp<-data.frame(dayinf=vector(),dayhosp=vector())
 if (sum(tohosptot)>0) {
     daysinfhosp<-hosptodayinf(daysinfhosp,tohosptot,time)

     # Also want to record the length of the time they spent being infectious, to see what the effect of
     # hospitalization is on infectious duration
     # If they were hospitalized on the same day as they became infectious, that means that their infectious
     # period was 1 day, so add 1 to the difference between dayhosp and dayinf
     inf_lengths<-c(inf_lengths,daysinfhosp$dayhosp-daysinfhosp$dayinf+1)
    }
    list(out1=x,out2=daysinfhosp,out3=inc_lengths,out4=inf_lengths, out5=onestep_Rdays)}

  #Data extraction functions
  #summary stats

  detcases<-function(firstday,lastday)
   #this counts only detected cases whose symptoms appear in the time window
  {sum((Zdays$dayinf>=firstday) & (Zdays$dayinf<=lastday))}

   detplusprevcases<-function (firstday,lastday) #detected cases -- those that are hospitalized
  (assume if you know about them you hospitalize them)
     #plus those left over in the I group on the last day of monitoring
  {Z$ht[min(84,lastday)]-Z$ht[firstday]+Z$it[min(84,lastday)]-Z$it[firstday]}

  detplusrecplusprevcases<-function (firstday,lastday) #the larger group those that were
hospitalized or recovered or prevalent(counted on the day of recovery)
  {detplusprevcases(firstday,min(84,lastday))+Z$rt[min(84,lastday)]-Z$rt[firstday]}

   checksum <-function(Z){(max(Z$nt)==min(Z$nt)) & min(Z$nt)==m+1}
   # END CORE FUNCTIONS
   botharm_totals<-vector()
   botharm_allcases<-vector()
   cat("Daily probability of case detection: ", pdhosp, "\n")
   cat("Maximum VES: ",VEmax, "\n")
   cat("Counting cases from day",startcount,"to day",endcount,"\n\n")
   # Initialize dataset to record survival times, censoring status, and trial arm. We’ll do a
 Cox PH model
   # with these data.
   eventtimes<-data.frame(eventday=vector(),eventstatus=vector(),treated=vector())

   initialI<-vector()
   initialE<-vector()
   IatVacc<-vector()
   EatVacc<-vector()
   Is_imm<-vector()
   Is_del<-vector()
   Es_imm<-vector()
   Es_del<-vector()

   for(ivac in 1:2){
     cat("\n")
     tvac<-tvacs[ivac]#set day of ring vaccination for this condition
     cat("tvac=",tvac,"\n")
     totals<-vector() firstmonth<-vector()
     allcases<-vector()

     incubation_periods<-vector()
     infectious_periods<-vector()

     numtrials<-0
     numsims<-0
     # Not every simulation will lead to a simulated trial, so only record those
     # that do, and go until we’ve done nsim simulated trials

     while (numtrials<nsim) {
         numsims<-numsims+1
         #initialize one ring simulation
         stateinit<-list(SU=m,SV=rep(0,nSV),EU=c(rep(0,maxnE-
 1),1),EV=rep(0,maxnE),IU=rep(0,maxnI),IV=rep(0,maxnI),RU=0,RV=0,H=0) #start with a single E
       time=-1
      dotrial<-0
      #set it going until a case is detected or until the epidemic runs out
      while (sum(stateinit$EU)+sum(stateinit$IU)>0) {
        list[stateinit]<-update(stateinit,pdhosppre)
        if (stateinit$H>0) {
          # if a case is detected, go ahead with the trial
           dotrial<-1
          numtrials<-numtrials+1
           break
        }
      }
     if (dotrial==1) {
       if (ivac==1) {
          initialI<-c(initialI,sum(stateinit$IU))
          initialE<-c(initialE,sum(stateinit$EU))} else {}
       if ((sum(stateinit$EU)+sum(stateinit$IU)==0) && (extfoi_numerator==0)) {

     totals<-c(totals,0)
     firstmonth<-c(firstmonth,0)

     allcases<-c(allcases,0)

     if (ivac==1) {
        IatVacc<-c(IatVacc,0)
        EatVacc<-c(EatVacc,0)
        Is_imm<-rbind(Is_imm,rep(0,endfu))
        Es_imm<-rbind(Es_imm,rep(0,endfu))
      } else {
        Is_del<-rbind(Is_del,rep(0,endfu))
        Es_del<-rbind(Es_del,rep(0,endfu))
      }
    } else {

      newsusceptibles=m+1-sum(stateinit$EU)-sum(stateinit$IU)
state<-
list(SU=newsusceptibles,SV=rep(0,nSV),EU=stateinit$EU,EV=rep(0,maxnE),IU=stateinit$IU,IV=rep(0,maxnI),RU=0,RV=0,H=0) #start with a single E
       time=1
       st<-rep(0,endfu)
       et<-rep(0,endfu)
       it<-rep(0,endfu)
       rt<-rep(0,endfu)
       ht<-rep(0,endfu)
       nt<-rep(0,endfu)
       Z<-data.frame(st,et,it,rt,ht,nt)

       Zdays<-data.frame(dayinf=vector(),dayhosp=vector()) #records day of start of
infectious period for all cases
       Rdays<-c() # records day on which each subject who moved into R (if they did)
        onesiminflength<-vector()
        onesiminclength<-vector()
        inclengths<-vector()
        inflengths<-vector()
        daysinfhosp<-vector()
        onestep_Rdays<-c()

        for (time in 1:endfu){ #CENTRAL LOOP IN ONE SIMULATION
           Z$st[time]<-sum(state$SU)+sum(state$SV)
           Z$et[time]<-sum(state$EU)+sum(state$EV)
           Z$it[time]<-sum(state$IU)+sum(state$IV)
           Z$ht[time]<-state$H
           Z$rt[time]<-state$RU+state$RV Z$nt[time]<-
     sum(state$SU)+sum(state$SV)+sum(state$EU)+sum(state$EV)+sum(state$IU)+sum(state$IV)+state$RU+s tate$RV+state$H
           onesiminclength<-c(onesiminclength,inclengths)
           onesiminflength<-c(onesiminflength,inflengths)
           Zdays<-rbind(Zdays,daysinfhosp)
           Rdays<-c(Rdays,onestep_Rdays)

           list[state,daysinfhosp,inclengths,inflengths,onestep_Rdays]<-update(state,pdhosp)
    }
      if (ivac==1) {
       # Record how many I and E there were at vaccination
      IatVacc<-c(IatVacc,Z$it[tvac])
      EatVacc<-c(EatVacc,Z$et[tvac])

      # Record the trajectory of Is and Es overall
      Is_imm<-rbind(Is_imm,Z$it)
      Es_imm<-rbind(Es_imm,Z$et)} else {
          Is_del<-rbind(Is_del,Z$it)
         Es_del<-rbind(Es_del,Z$et)
       }
    # Add information to the survival data set for cases
    # Specifically, survival times are hospitalization times from Zdays,
    # event status is 1 if they occur in the time window,
    # and treatment arm is 2-ivac (1 if immediate, 0 if delayed)
    cases<-Zdays[(Zdays$dayinf>=startcount) & (Zdays$dayinf<=endcount),]
    num_cases<-length(cases$dayinf)
    eventtimes_cases<-data.frame(eventday=cases$dayinf,
                                                eventstatus=rep(1,num_cases),
                                                treated=rep(2-ivac,num_cases))
    recovered<-Rdays[(Rdays>=startcount) & (Rdays<=endcount)]
    num_recovered<-length(recovered)
    eventtimes_recovered<-data.frame(eventday=recovered,
                                                eventstatus=rep(0,num_recovered),
                                                treated=rep(2-ivac,num_recovered))
    # The number in the risk set at t=startcount is all individuals (m+1), minus those
  have already
    # been recovered or hospitalized at that stage. Those who are E or I are technically
  not
    # in the risk set, but we can’t tell if they are E or I until we detect them.
  Therefore,
    # include them, and if we detect them later we can either class them as an event (if
symptoms
    # developed in the window) or censored (if they developed before the window)
    num_riskset<-m+1-Z$rt[startcount]-Z$ht[startcount]
    num_censored<-num_riskset-num_cases-num_recovered
    eventtimes_censored<-data.frame(eventday=rep(endcount,num_censored),
                                                eventstatus=rep(0,num_censored),
                                                treated=rep(2-ivac,num_censored))
    eventtimes<-
rbind(eventtimes,eventtimes_cases,eventtimes_recovered,eventtimes_censored)

      incubation_periods<-c(incubation_periods,onesiminclength)
      infectious_periods<-c(infectious_periods,onesiminflength)

      totals<-c(totals,detcases(startcount,endcount))
      firstmonth<-c(firstmonth,detcases(1,31))

      allcases<-c(allcases,detplusrecplusprevcases(startcount,endcount))
     }
   } else {}
  }
  botharm_totals<-c(botharm_totals,totals)
  botharm_allcases<-c(botharm_allcases,allcases)

  SAR[ivac]<-mean(totals)/m
  trueSAR[ivac]<-mean(allcases)/m
  cat("detected cases first month",mean(firstmonth)*100/m,"%\n") #detected SAR in first
month
  cat("detected cases during followup",SAR[ivac]*100,"%\n") #detected SAR in followup
  cat("all cases during followup",trueSAR[ivac]*100,"%\n") #actual SAR
 }
 # Sample size using a binomial test for proportions
ssu<-samplesize.bin(.025,.8,SAR[1],SAR[2])/m # Outputs total sample size
truessu<-samplesize.bin(.025,.8,trueSAR[1],trueSAR[2])/m

 # Sample size using a Cox’s proportional hazards model. See code updates document for assumptions
 survmodel<-coxph(Surv(eventday,eventstatus)~treated,eventtimes)
 # Make a new data set with one treated and one untreated
 newdata<-data.frame(eventdays=c(0,0),eventstatus=c(0,0),treated=c(0,1))
 # Get predicted values for each
 predicted<-predict(survmodel,newdata)
 # Calculate the hazard ratio from these predicted values
 HR<-exp(predicted[2]-predicted[1])
 # Sample size calculation (formula is from Hsieh and Lavori 2000)
 surv_deaths<-4*(1.96 + 0.84)^2 / (log(HR)^2)
 surv_ssu<-surv_deaths/(mean(SAR)*50)
 # ICC calculation with covariance adjustment is from Shoukri et al botharm_totals_sq<-botharm_totals^2
 K<-2*nsim
 # Covariance-adjusted ANOVA estimator of ICC
 MSB<-1/(K-1) * (sum(botharm_totals_sq)/m - sum(botharm_totals)^2/(m*K))
 MSW<-1/(K*(m-1)-1) * (sum(botharm_totals) - sum(botharm_totals_sq)/m)
 ICC_detected<-(MSB-MSW)/(MSB+(m*(K-2)/(K-1)-1)*MSW)
 deff_detected<-1+(m-1)*ICC_detected
 # ICC calculation for true cases botharm_allcases_sq<-botharm_allcases^2
 # Covariance-adjusted ANOVA estimator of ICC for true cases
 MSB<-1/(K-1) * (sum(botharm_allcases_sq)/m - sum(botharm_allcases)^2/(m*K))
 MSW<-1/(K*(m-1)-1) * (sum(botharm_allcases) - sum(botharm_allcases_sq)/m) ICC_true<     -(MSB-MSW)/(MSB+(m*(K-2)/(K-1)-1)*MSW)
 deff_true<-1+(m-1)*ICC_true
 # Make the confidence interval for the SAR difference
 SARdiff<-SAR[1]-SAR[2]
 SARbar<-(SAR[1]+SAR[2])/2
 varSARdiff<-SARbar*(1-SARbar)*(2/(nsim*m))*deff_detected
 SARdiff_ci_lower<-SARdiff-1.96*sqrt(varSARdiff)
 SARdiff_ci_upper<-SARdiff+1.96*sqrt(varSARdiff)

 VEest<-(1-SAR[1]/SAR[2])*100
 sd_ve<-sqrt((1/(SAR[1]*m*nsim) + 1/(SAR[2]*m*nsim) - 2/(nsim*m))*deff_detected)
 ci_lower<-1-(1-VEest/100)*exp(1.96*sd_ve)
 ci_upper<-1-(1-VEest/100)*exp(-1.96*sd_ve)

 trueVEest<-(1-trueSAR[1]/trueSAR[2])*100
 truesd_ve<-sqrt((1/(trueSAR[1]*m*nsim) + 1/(trueSAR[2]*m*nsim) - 2/  (nsim*m))*deff_true)
 trueci_lower<-1-(1-trueVEest/100)*exp(1.96*truesd_ve)
 trueci_upper<-1-(1-trueVEest/100)*exp(-1.96*truesd_ve)

 cat("\n Estimated efficacy based on SAR: ",VEest,"%\n")
 cat ("Total sample size uncorrected for clustering: ",ssu," rings of ",m,"persons\n")
 cat("sample size corrected (approximately)",deff_detected*ssu,"\n")
 cat("Approximate CI for VE: (",ci_lower,",",ci_upper,")\n")
 cat("Approximate CI for SAR difference: (",SARdiff_ci_lower,",",SARdiff_ci_upper,")\n")

list(out1=VEest,out2=ci_lower,out3=ci_upper,out4=ssu,out5=deff_detected,out6=SAR,out7=ICC_dete cted,
      out8=trueVEest,out9=trueci_lower,out10=trueci_upper,out11=truessu,out12=deff_true,
      out13=trueSAR,out14=ICC_true,out15=surv_deaths,out16=initialE,out17=initialI,out18=Es_imm,out1
       9=Is_imm,out20=Es_del,out21=Is_del)
}
~~~

